# Spectrolipidomics of glial cell lines: a deuterated probe for semiquantitative monitoring of cannabidiol-induced cholesterol modulation

**DOI:** 10.64898/2026.01.26.701843

**Authors:** K. Chrabąszcz, T. Kossowski-Kołodziej, A. Panek, K. Pogoda

## Abstract

Understanding lipid metabolism in peripheral glial cells is crucial for elucidating the molecular mechanisms underlying neurodegeneration, cancerogenesis and therapy resistance. Here, we introduce a spectrolipidomic sensing approach that integrates Raman, FT-IR, and AFM-IR spectroscopy to monitor nanoscale cholesterol remodeling in glial cells exposed to cannabidiol (CBD). Deuterated cholesterol (dChol) was employed as an intrinsic, spectroscopically active molecular probe, enabling selective tracking of cholesterol transformations through characteristic C–D vibrational signatures within the 2300–2000 cm^−1^ silent spectral region. Multimodal vibrational spectroscopy provided label-free, spatially resolved insight into lipid organization, redistribution, and metabolic reprogramming across micro- and nanoscales. The dChol probe enabled semi-quantitative evaluation of cholesterol uptake, esterification, and membrane integration, revealing that the sequence of CBD exposure, before or after probe addition, triggers distinct lipid metabolic pathways. Raman spectroscopy demonstrated superior sensitivity, with reliable detection of intracellular dChol at concentrations as low as 10 µM, outperforming FT-IR imaging and confirming its suitability for cell lipid sensing. This analytical platform establishes deuterium-labeled lipids as powerful vibrational sensors for probing lipid metabolism and CBD-induced remodeling in situ. The presented spectrolipidomic framework paves the way for next-generation, spectroscopy-based biosensing systems capable of visualizing lipid dynamics, membrane restructuring, and drug– lipid interactions under pharmacological or environmental stress conditions.

**Highlights:**

- Deuterated cholesterol (dChol) used as an intrinsic vibrational sensor
- Lower detection threshold of intracellular dChol for Raman than FT-IR
- AFM-IR reveals phases of lipid droplet formation in nanoscale
- CBD alters cholesterol uptake, esterification, and lipid unsaturation profiles

## 1. Introduction

Lipids play a fundamental role in cellular physiology, forming the structural and functional basis of biological membranes while participating in energy storage, signaling, and stress response mechanisms. In peripheral glial cells, including Schwann cells and malignant peripheral nerve sheath tumor (MPNST) cells, lipid composition is highly dynamic and reflects both the specialized functions of glial membranes in nerve regeneration and the biochemical reprogramming associated with tumorigenic transformation.[1],[2],[3] Moreover, in MPNST, one of the most aggressive soft tissue sarcomas of the nervous system, deregulated lipid metabolism contributes to uncontrolled proliferation, invasiveness, and therapy resistance.[4]

Among bioactive compounds influencing lipid metabolism, cannabidiol (CBD) has gained significant attention due to its regulatory effects on cellular homeostasis, oxidative stress, and membrane organization.[5],[6] CBD modulates lipid pathways by interacting with the endocannabinoid system and enzymes involved in lipid synthesis and degradation.[7] Several studies have demonstrated that CBD alters cholesterol metabolism, lipid droplet formation, and membrane fluidity, thereby influencing signaling, vesicle trafficking, and mitochondrial activity.[8],[9],[10],[11] Moreover, membrane lipid composition, particularly cholesterol content and lipid order, can modulate cellular responses to anticancer therapies, including radiotherapy.[12],[13],[14] Alterations in membrane rigidity, redox status, and lipid raft structure affect the generation of reactive oxygen species (ROS) and the activation of death pathways following ionizing radiation. [15],[16] Lipid peroxidation also underlies ferroptosis, an iron-dependent cell death process triggered by excessive lipid oxidation. [17],[18] Consequently, modulation of lipid composition by compounds such as CBD may influence radiosensitivity by regulating ROS levels and lipid peroxidation. Despite these insights, the direct biochemical consequences of CBD-induced lipid remodeling remain poorly understood, largely due to the lack of non-destructive analytical tools capable of visualizing lipid transformations in situ.

To address these analytical challenges, vibrational spectroscopy offers a powerful, label-free approach for studying lipid organization, dynamics, and biochemical remodeling within intact cells.[19],[20],[21] Raman and infrared (IR) spectroscopy provide unique spectral fingerprints that allow selective tracking and semiquantitative analysis of lipid composition and distribution under various conditions. Here, we introduce the concept of spectrolipidomics, the application of vibrational spectroscopy combined with isotopically labeled lipid probes to study lipid metabolism and organization in living cells. This approach offers a chemically specific, non-invasive framework for monitoring lipid transformations and correlating molecular composition with cellular function.

Deuterated lipid analogs serve as intrinsic spectroscopic probes, introducing characteristic C–D stretching vibrations in the 2300-2000 cm^−1^ “silent” spectral window, where endogenous biomolecules contribute minimally.[22],[23],[24] Their incorporation allows selective detection of lipid localization and dynamics within complex cellular environments. Similar principles have been applied using azide probes as infrared metabolic sensors.[25] In contrast to conventional methods such as mass spectrometry (MS) or nuclear magnetic resonance (NMR), which require destructive sample processing, vibrational spectroscopy enables non-invasive, spatially resolved biochemical analysis of intact systems.

In this study, we propose a spectroscopic approach combining Raman, FT-IR and AFM-IR analyses to investigate CBD-induced lipid remodeling in peripheral glial cells. Using deuterated cholesterol as a spectroscopic probe, we establish a novel sensing strategy for semi-quantitative tracking of cholesterol dynamics through characteristic C–D vibrational modes. Nanoscale analysis relied on alternative lipid-associated vibrational markers, including the ester carbonyl band at ∼1740 cm^−1^, to indirectly monitor the incorporation and redistribution of dChol within cellular lipid structures. This spectrolipidomic framework provides a label-free, chemically specific tool for probing drug–lipid interactions and metabolic responses in native cellular environments, advancing molecular sensing in lipid biology and cancer research.

## 2. Materials and Methods

### 2.1. Reagents

#### Cannabidiol

A stock solution of cannabidiol (CBD; Merck) was purchased as 1.0 mg/mL solution in methanol and stored at −20 °C, according to manufacturer ‘s recommendation. The stock was serially diluted in methanol to obtain 1000 µM concentration. For cell treatment, the 1000 µM solution was further diluted with culture medium to reach a final CBD concentration of 3 µM. The methanol content in all working solutions was adjusted to remain constant at 0.001% (v/v), ensuring that the solvent had no measurable influence on cellular biochemistry. Control samples were treated with an equivalent volume of methanol (0.001%) to maintain experimental consistency and ensure reliable comparison between CBD-treated and control cells.

#### Cholesterol and deuterated cholesterol

Methyl-*β*-cyclodextrin, cholesterol and d_6_-cholesterol were purchased from Sigma. Cholesterol and d_6_-cholesterol/methyl-*β*-cyclodextrin stock complexes were prepared using a methyl-*β*-cyclodextrin to cholesterol (d_6_-cholesterol) ratio of 6.2:1 [22] and a final concentration of 1µM, 10µM, 50µM and 100µM for both compounds. The various concentrations of cholesterol were implemented only for the comparison of possible H^2^ cytotoxic effect from d_6_-cholesterol on studied cells investigated by MTS viability assay (Supporting Information, Fig. S1).

### 2.2. Cell Culture, CBD and Lipid Administrations, MTS assay

#### Cell culture

Human Schwann (normal, hTERT NF1 ipnNF95.11c) and malignant MPNST (cancer, sNF02.2) cell lines were purchased and cultured according to ATCC protocol in DMEM medium, supplemented with 10% of FBS (fetal bovine serum), 100 U/ml penicillin– streptomycin solution. MPNST cell line was additionally supplemented with L-glutamine. The cells were grown in 75 cm^3^ flasks at 37°C in an atmosphere of 5% CO_2_. Prior to each experimental treatment, cells were serum-starved for 2 h in Dulbecco’s phosphate-buffered saline (DPBS) containing Ca^2+^ and Mg^2+^. Following serum starvation, cells were subjected to cholesterol or d_6_-cholesterol treatments at final concentrations of 1, 10, 50, or 100 µM. To investigate the order-dependent effects of cannabidiol (CBD) and d_6_-cholesterol, two sequential incubation protocols were applied. In the first protocol, cells were incubated with CBD for 24 h, after which d_6_-cholesterol was added and incubation was continued for an additional 24 h. In the second protocol, cells were first incubated with d_6_-cholesterol for 24 h, followed by CBD treatment for a further 24 h.

The experimental workflow was visualized on Fig. 1.

**Figure 1.**
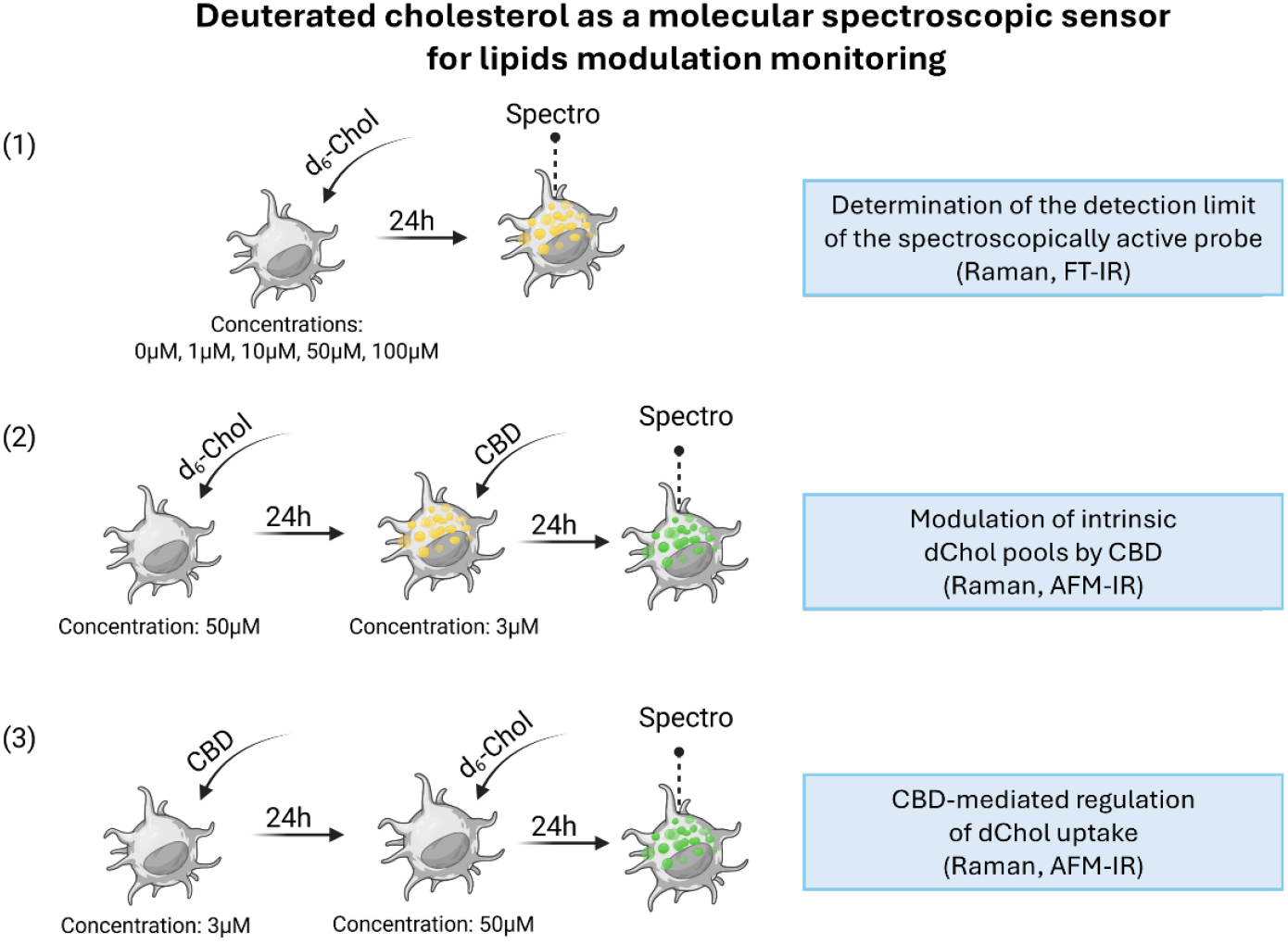
Conceptual overview of the spectrolipidomic sensing platform employing deuterated cholesterol probes for semiquantitative Raman, FT-IR and AFM-IR detection of lipid modulation.

#### CBD incubation

A CBD concentration of 3 µM was selected based on its previously demonstrated effects on lipid metabolism, as reported in our earlier studies.[11] Both cell lines were incubated with 3µM CBD concentration 24h before or after d_6_-cholesterol administration.

#### Cholesterol/d_6_-cholesterol administration

The prepared solutions of cholesterol/cyclodextrin and d_6_-cholesterol/cyclodextrin with 1µM, 10µM, 50µM and 100µM concentrations were applied to cell culture. The use of this concentration range for d_6_-cholesterol enabled determination of the limit of detection for the spectroscopic analyses. Based on these results, a concentration of 50 µM d_6_-cholesterol was selected for subsequent experiments in combination with CBD. Moreover, the Raman and FT-IR spectra of cholesterol and d_6_-cholesterol are presented at Fig. S2 (Supporting Information).

#### MTS assay

CBD influence on the cells viability and metabolic activity was already investigated in our previous studies.[11] For lipids cytotoxicity the cells were tested using CellTiter 96 AQueous One Solution Cell Proliferation Assay (Promega) with tetrazolium compound. Both cell lines were investigated after 24 h of incubation with 1µM, 10µM, 50µM and 100µM concentration of cholesterol/cyclodextrin (d_6_-cholesterol/cyclodextrin) and after 48h for the mixtures of 50µM d_6_-cholesterol/3µM CBD and 3µM CBD/50µM d_6_-cholesterol.

Cells were seeded in 24-well plates at a density of 15,000 /well and 24h after seeding cells were serum-starved for 2h in DPBS with Ca^2+^ and Mg^2+^. Next, cells were treated with a selected concentration of implemented compounds. CellTiter 96® AQueous One Solution reagent was added directly to the culture wells and incubated for 1 hour at 37°C. Then the absorbance of formazan was recorded at 490 nm with a Spark 10 M (Tecan) multimode microplate reader. The absorbance values obtained with the untreated cells (control, additionally treated with methanol and methyl-*β*-cyclodextrin) were used for data normalization. Assays were conducted in triplicate. The MTS assays results can be found in Supporting Information, Fig. S1.

### 2.3. Spectroscopic investigations

#### Raman, FT-IR, and AFM-IR measurements

All spectroscopic measurements were performed on cells seeded on calcium fluoride windows (Crystran Ltd., UK) inside 12-well plates and kept in an incubator at 5% CO2 and 37°C for 24 h to promote adhesion and growth. Then particular set of samples were treated with (1) 1µM, 10µM, 50µM and 100µM concentration of d_6_-cholesterol/cyclodextrin to determine the spectroscopic limit of detection, (2) 50µM d_6_-cholesterol, (3) 3µM CBD. After 24 hours (1) set of samples were rinsed twice with phosphate-buffered saline (PBS) for 5 min and fixed with 4% paraformaldehyde (PFA) in PBS for 20 min. Then, all samples were washed with PBS (3 times for 2 min) to remove PFA residues. (2) were incubated with 3µM CBD, (3) were incubated with 50µM d_6_-cholesterol and left for 24 hours (Fig. 1). Then also these samples (2 and 3) were fixed with PFA as described above. For Raman measurements, samples were left in PBS solution. Before FT-IR and AFM-IR measurements samples were washed twice in ultrapure water and then dried under a gentle stream of nitrogen. This sample preparation protocol was selected based on our recently published work, which demonstrated that it provides optimal preservation of both molecular composition and morphological integrity of the samples.[26] All solutions were prepared using ultrapure water (DirectQ 3 UV, Millipore, USA).

Raman imaging of individual cells was performed using a Renishaw InVia Raman spectrometer integrated with a confocal optical microscope. Excitation was provided by an air-cooled solid-state laser (λ = 532 nm), and the scattered light was detected with a CCD detector maintained at −70 °C. A 60× water-immersion objective (LUMPlanFL, NA 1.0; Olympus) was applied to achieve high spatial resolution and minimize refractive index mismatches during imaging. Raman maps were collected from the entire cell area for 15 cells per condition, using a 0.5 µm step size, an integration time of 0.3 s (1 accumulation), and a spectral resolution of approximately 1.5 cm^−1^. The spectrometer calibration was performed using an internal silicon standard. Single-cell FT-IR hyperspectral images were acquired using a HYPERION 3000 infrared microscope (Bruker, Ettlingen, Germany) equipped with a 36× objective and coupled to a VERTEX 70v spectrometer operating in transmission mode. The focal plane array (FPA) detector captured images with a 64 × 64-pixel format, corresponding to a projected pixel size of 1.1 µm × 1.1 µm, enabling spatially resolved chemical mapping of cellular components. FT-IR images were acquired from 15 cells for each condition within the spectral range of 900–3800 cm^−1^, at a spectral resolution of 4 cm^−1^ and 256 scans per spectrum. AFM-IR nanospectroscopy investigations were performed using a NanoIR2 spectrometer (Anasys Instruments, Santa Barbara, CA, USA) operating in contact mode. Measurements were conducted with silicon gold-coated PR-EX-nIR2 probes (tip diameter: ∼30 nm; resonance frequency: 13 ± 4 kHz; Anasys Instruments, USA). Contact resonances were selected using an 180 kHz search location and a 50 kHz half-width Gaussian filter. Spectra were collected using a multichip tunable quantum cascade laser (QCL; MIRcat-QT, Daylight Solutions) covering the 3000–2800 cm^−1^ and 1800–900 cm^−1^ ranges, with a spectral resolution of 2 cm^−1^. For each condition, spectra were recorded from 10 points positioned diagonally across 10 individual cells, by co-averaging 256 excitation pulses. For AFM topography acquisition, the cantilever scan rate was set to 0.05 Hz. The spatial resolution was approximately 160 nm, as the measured area (80 × 80 µm) comprised 500 × 500 measurement points.

#### Data analysis

Raman images were processed using WiRE software (ver.5.3, Renishaw, United Kingdom). The steps included removal of cosmic rays, noise filtering, baseline correction, and min-max normalization of all spectra. Subsequently, hierarchical cluster analysis (HCA) was applied to distinguish subcellular components based on their spectral profiles. Euclidean distance and Ward’s method were used to determine spectral distances and define individual clusters.

FT-IR data were preprocessed using Matlab 2017 and OPUS software (ver. 7.5, Bruker Optics, Germany). Mean spectra were extracted from individual cells, followed by min-max normalization in the amide I region to minimize effects related to sample thickness. The second derivative of the IR spectra was then calculated using the Savitzky–Golay method with 13 smoothing points.[11]

AFM-IR spectra were analyzed with Analysis Studio (ver. 3.14). Spectra were smoothed using a third-order Savitzky–Golay polynomial with five data points, normalized via min-max normalization, and presented as second derivative spectra.[27] Three-dimensional AFM and AFM-IR images were generated using MountainsMap software (ver. 7.3, Digital Surf, France).

The areas under selected Raman, FT-IR, and AFM-IR bands (integral intensities) were calculated using OPUS (ver. 7.5, Bruker Optics, Germany). Box plots were then constructed from these values to visualize biochemical variations within cells. Statistical analysis of variance was performed using ANOVA in OriginPro software, and Tukey’s test was applied to determine significance levels (p-values).

Principal Component Analysis (PCA) was performed using Unscrambler X (ver. 10.3). Depending on the dataset, PCA was calculated for selected spectral regions using a leave-one-out cross-validation method and the Nonlinear Iterative Partial Least Squares (NIPALS) algorithm for data decomposition. The initial decomposition included seven principal components (20 iterations, NIPALS algorithm), and the results were visualized as three-dimensional score plots accompanied by corresponding loading plots.

## 3. Results and Discussion

### Evaluating detection limits of deuterated probe via Raman and FT-IR spectrolipidomics

The limit of detection (LOD) for the proposed deuterated probe (d_6_-cholesterol, dChol) was determined using both Raman and FT-IR spectroscopy (Fig. 2). Following administration dChol at initial concentrations of 1-100 µM to peripheral glial cell lines, Raman spectroscopy enabled tracking of probe accumulation within the cytoplasm based on C–D band integration in hyperspectral maps (Fig. 2, C-E). Owing to the capability of Raman spectroscopy to measure cells in buffer, thus preserving their native morphology and volume, well-defined lipid droplets (LDs) of various sizes were clearly visualized. At higher probe concentrations (≥10 µM), LD accumulation increased markedly around the nucleus. Cluster analysis differentiated the mean spectra of LDs, cytoplasm, and nuclei, confirming LD localization within the cytoplasmic region without affecting nuclear composition (Fig. S3, C-D).

**Figure 2.**
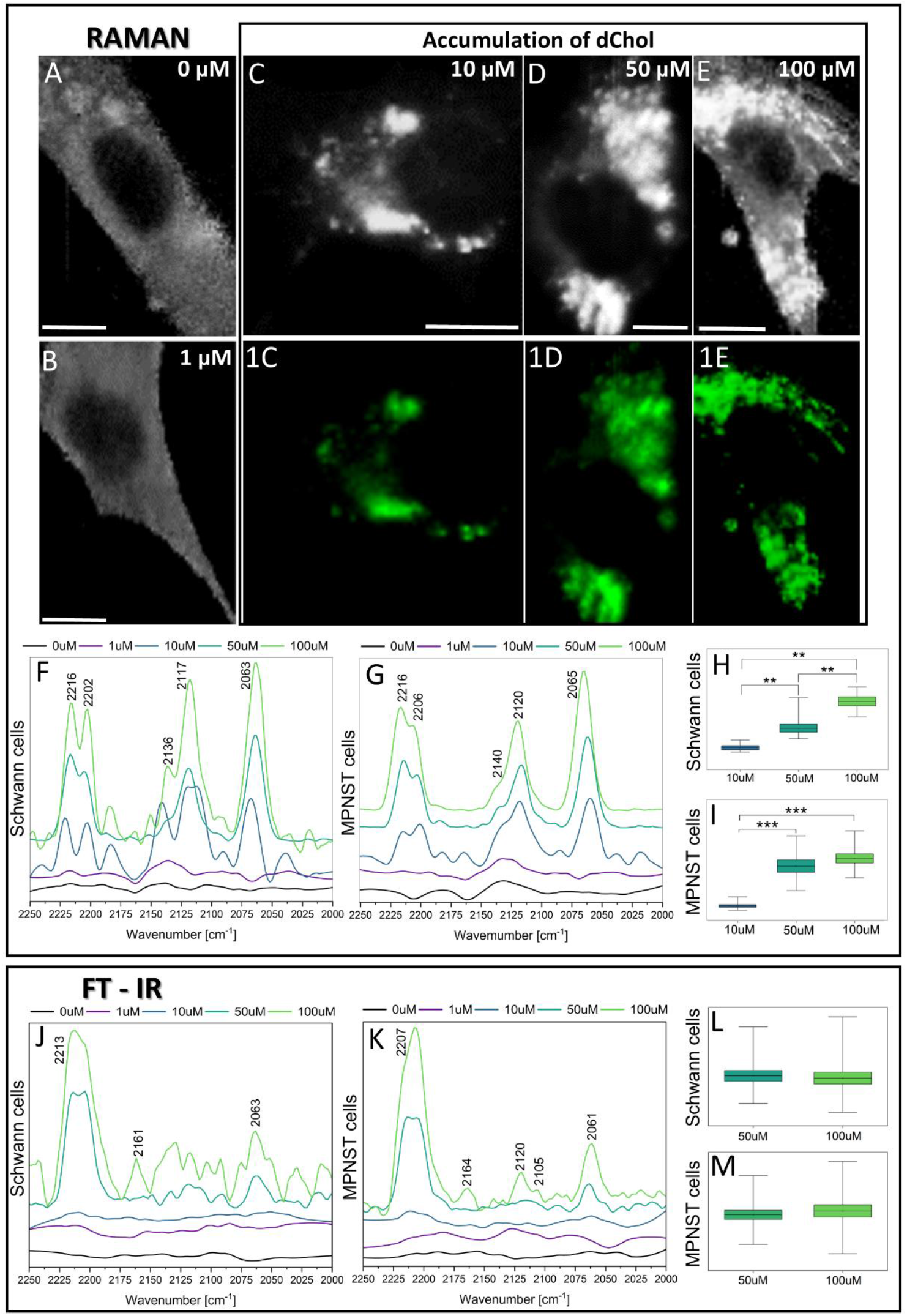
Exemplary chemical Raman maps of peripheral glia cells body (A-E) and distribution of C– D bands within accumulated lipid droplets (1C-1E) following incubation with 0 µM, 1 µM, 10 µM, 50 µM, and 100 µM concentrations of d_6_-cholesterol. Scale bar equals 10µm. Spectral ranges corresponding to the cytoplasm with assigned C–D vibrational bands are shown for Raman (F, G) and FT-IR (J, K) spectroscopy. The sum of integrated C–D bands intensities calculation for Raman (H, I) and FT-IR (L, M) spectra is also presented. Statistical significance was determined using the one-way ANOVA (***p<0.001, p**<0.01).

Based on mean cytoplasmic spectra, the LOD for dChol was evaluated for both cell lines (Fig. 2, F, G). The characteristic C–D bands appeared between 2250 and 2000 cm^−1^ (at 2216 cm^−1^, 2120 cm^−1^, and 2065 cm^−1^), consistent with marker bands observed in the pure dChol spectrum (Fig. S2, A). For both cell types, these bands became clearly detectable at concentrations ≥10 µM, with intensity increasing in a dose-dependent manner. Integration of the C–D bands enabled semi-quantification of intracellular probe levels. In Schwann cells, the correlation between deuterium-related band intensity and concentration was linear (Fig. S4, A. R^2^ = 0.9992), indicating consistent uptake and distribution. In contrast, MPNST cells showed weaker correlation (Fig. S4, B. R^2^ = 0.8212), suggesting greater signal heterogeneity likely linked to altered lipid metabolism or structural organization in the cancer phenotype (Fig. 2, H, I). [28],[29]

The same experiment performed using FT-IR imaging required sample drying due to water absorption overlapping with cellular spectra. Despite this limitation, C–D bands remained detectable (Fig. 2, J, K), though only at higher probe concentrations (50-100 µM). The observed pattern was consistent with the pure dChol spectrum (Fig. S2, B). However, integrated band intensity analysis revealed no statistically significant difference between the 50 µM and 100 µM samples (Fig. 2, L, M). Therefore, Raman spectroscopy proved more suitable for semi-quantitative evaluation of active deuterated lipid probes in cells.

#### Tracing deuterated cholesterol uptake in glial cells beyond C–D vibrations at nanoscale

High-resolution AFM imaging of Schwann and MPNST cells revealed distinct topographical alterations following dChol probe administration (Fig. 3, A-J). During sample drying, LDs lost their spherical shape seen in Raman imaging, leaving voids in the cytoplasm corresponding to their original locations (Fig. 3, D-E, I-J, yellow arrows). Height profiles across the nuclei (Fig. 3, 1A-E, red traces) showed that increasing dChol concentration did not affect nuclear height in Schwann cells, which remained around 800 nm. In contrast, MPNST cells displayed marked nuclear remodeling: while low probe levels (1-10 µM) remain similar heights (∼800 nm), greater concentrations (50-100 µM) caused pronounced deformation and elevation, reaching ∼1600 nm (Fig. 3, 1J). Cytoplasmic thickness analysis showed that untreated Schwann cells had approximately fourfold thicker cytoplasm than MPNST cells (100 nm vs. 400 nm; Fig. 3, 2A, 2F). At greater probe concentrations (50-100 µM), both cell lines exhibited an additional increase of 200-500 nm in Schwann and 300-800 nm in MPNST cells. Remarkably, the cavities left after LD assembly in MPNST cells treated with 100 µM dChol reached ∼1400 nm in depth. These differences likely reflect metabolic reprogramming typical of cancer cells. [29] Schwann cells, with slower lipid turnover, incorporated dChol without excessive accumulation, as indicated by larger cytoplasmic signal variation between concentrations (Fig. 2, H). Conversely, MPNST cells, characterized by elevated lipid demand, displayed greater probe uptake and smaller inter-concentration differences due to faster lipid cycling (Fig. 2, I).[30] Such lipid remodeling may also influence nuclear architecture, consistent with previous reports linking LDs expansion to nuclear indentation in oleic acid treated hepatic cells.[31]

**Figure 3.**
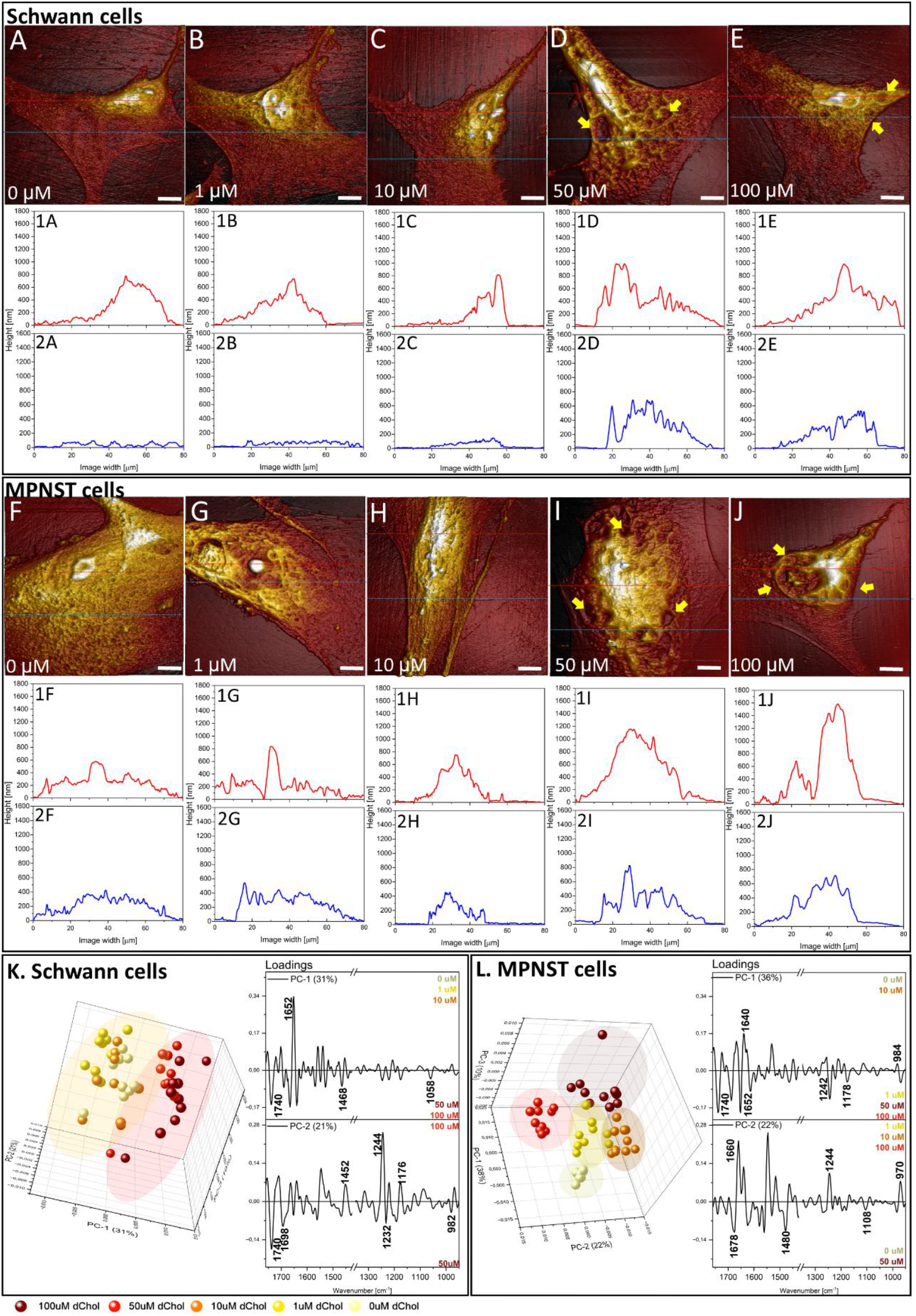
Representation of 3D AFM topography maps for Schwann (A-E) and MPNST (F-J) treated with 0 µM, 1 µM, 10 µM, 50 µM, and 100 µM concentrations of d_6_-cholesterol. Scale bar equals 10µm. Height profiles correspond to cross-sections through the nuclear (red trace) and cytoplasmic (blue trace) regions were presented (1A-1E). Principal component analysis (PCA) was applied to AFM-IR spectra to identify factors associated with d_6_-cholesterol treatment beyond conventional C–D vibration region (K, L).

As AFM-IR employ segmented (QCL) infrared sources, such systems typically exclude the 2800-1800 cm^−1^ “silent” spectral region. Consequently, identification of deuterated probe incorporation by AFM-IR required alternative spectral markers. Principal component analysis (PCA) was therefore employed to identify non-C–D spectral features associated with dChol uptake (Fig. 3, K-L). In Schwann cells, PCA revealed clear separation between low (0-10 µM) and high (50-100 µM) probe concentrations in the 3D score plot (Fig. 3, K). High-concentration samples showed positive PC-1 scores linked to negative loadings, consistent with the second derivative preprocessing. An intensity increase at 1740 cm^−1^ (ester C=O stretch) with rising probe concentration suggested partial conversion of d_6_-cholesterol to d_6_-cholesteryl esters, preferentially stored in LDs.[32] At 100 µM, a more prominent change was observed at 1244 cm^−1^, attributed to nucleic acids, likely reflecting chromatin condensation due to LD-induced nuclear compression.[31] In MPNST cells, PCA results showed tighter clustering among conditions, indicating subtler spectral differentiation (Fig. 3, L). Nevertheless, higher probe concentrations correlated with increased 1740 cm^−1^ signal (negative PC-1 loading) and enhanced contribution at 1244 cm^−1^ (positive PC-2), suggesting a comparable, but less pronounced, spectral trend to Schwann cells.

Together, these findings highlight that deuterated cholesterol uptake in glial cells can be effectively monitored beyond conventional C–D vibration analysis. The combined AFM-IR and PCA approach revealed distinct lipid remodeling patterns associated with cellular phenotype and metabolic activity.

#### CBD influence on intracellular cholesterol modifications and uptake

To evaluate the effect of CBD on cholesterol dynamics, Schwann and MPNST cells were treated with dChol and CBD in two experimental configurations, as previously described (Fig. 1, 2-3). Representative LD visualizations based on C–D vibrations showed that, regardless of treatment sequence, Schwann cells formed markedly larger LDs than MPNST cells (Fig. 4, 1A-D). Using this spectroscopically active probe, we quantified CBD-dependent alterations in cholesterol handling and lipid-related parameters, including total lipid content, degree of unsaturation, and esterification level (Fig. 4, E-H).

**Figure 4.**
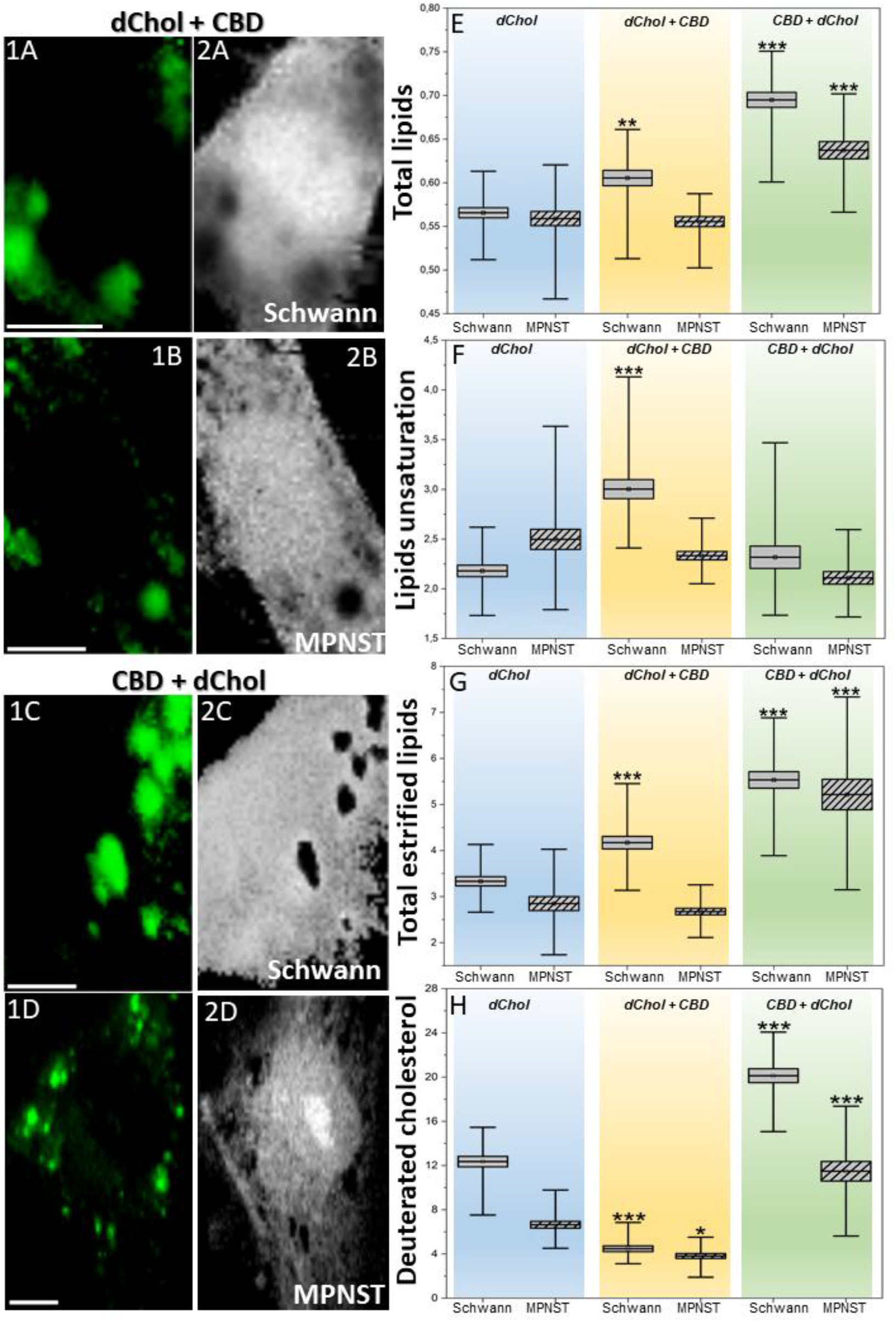
Distribution of d_6_-cholesterol-rich lipid droplets (green, LDs) within Schwann and MPNST cells treated with 50 µM d_6_-cholesterol/3 µM CBD (A, B) and 3 µM CBD/50 µM d_6_-cholesterol (C, D). Box diagrams (E-H) were generated from integrated intensity values obtained from mean spectra of lipid droplets, enabling semi-quantitative assessment of total lipids (E: 2850 cm^-1^/2930 cm^-1^), lipids unsaturation (F:1650 cm^-1^/1440 cm^-1^), total esterified lipids (G: 1740 cm^-1^/1000 cm^-1^) and deuterated cholesterol (H: (2220 cm^-1^+2120 cm^-1^+2065+cm^-1^)/1000 cm^-1^). Statistical significance was calculated using the one-way ANOVA (***p<0.001, p**<0.01, p*<0.05). Scale bar equals 10µm.

Following dChol administration, the overall LD accumulation was comparable between Schwann and MPNST cells (Fig. 4, E; mean values 0.57 and 0.58, respectively). Notably, CBD treatment applied after dChol uptake increased total lipid content only in Schwann cells (mean 0.61). In contrast, CBD exposure before dChol addition significantly enhanced probe accumulation in both cell types (Schwann: 0.69; MPNST: 0.64). The baseline degree of lipid unsaturation was similar in both lines (Fig. 4, F; Schwann 2.18; MPNST 2.50). When CBD was administered after probe uptake, Schwann cells exhibited a shift toward more unsaturated lipid species, while MPNST cells showed no such effect. Conversely, pre-treatment with CBD produced no measurable changes, suggesting that CBD promotes lipid desaturation primarily in already stored lipid pools. This indicates that CBD may enhance the synthesis or remodeling of unsaturated lipids in non-malignant cells. [33]

For esterified lipids, the predominant storage form within LDs (Fig. 4, G), CBD added after dChol increased esterification exclusively in Schwann cells, whereas CBD pre-treatment elevated esterified lipid levels in both cell lines. This conversion of free cholesterol to cholesteryl esters likely represents a protective mechanism limiting lipid peroxidation and enabling safe storage within LDs.[34] CBD is known to disturb cholesterol homeostasis by integrating into membranes and modulating enzymatic pathways involved in lipid esterification, droplet biogenesis, and sterol trafficking, thereby creating conditions favorable for enhanced cholesterol ester formation. [8],[35]

Analysis of the C–D spectral region provided further insight into dChol behavior under CBD exposure (Fig. 4, H). When CBD was applied after probe addition, dChol signal intensity decreased markedly in both cell lines (Schwann: 4.47; MPNST: 3.82), consistent with CBD-induced restriction of cholesterol accessibility due to altered membrane organization.[8] In contrast, when dChol was administered after CBD treatment, probe levels remained substantially higher (Schwann: 20.13; MPNST: 11.49), indicating that CBD reduces intracellular cholesterol only when lipid droplets are already formed.

#### AFM-IR nanospectroscopic analysis of CBD-induced lipid chain reorganization and morphological remodeling

To capture the broader biochemical responses occurring at the nanoscale, AFM-IR spectroscopy was employed. The mean second-derivative AFM-IR spectra (Fig. 5, A, B) for all investigated conditions revealed that the most pronounced spectral variations occurred within the protein region (1683-1636 cm^−1^).[11] Treatment with dChol alone or with subsequent CBD exposure did not alter the band positions or relative intensities in either the protein (1683-1636 cm^−1^) or phosphate (DNA/RNA/phospholipid; 1242-1220 cm^−1^) regions (Fig. 5, A, B blue and yellow traces). In contrast, when CBD was administered before dChol, a decrease in the 1636 cm^−1^ band was observed in both Schwann and MPNST cells, indicating a reduction in β-sheet protein components (Fig. 5, A, B green trace). [36] This observation agrees with previous findings that CBD attenuates amyloid aggregation and reduces β-sheet–rich conformations in amyloidogenic systems.[37] Interestingly, under the same condition, a new band at 1056 cm^−1^ appeared exclusively in Schwann cells, accompanied by an intensified 1220 cm^−1^ feature corresponding to the O–P–O moiety of RNA or phosphate esters (Fig. 5, A green trace).[36] The CBD-induced emergence of this signal suggests localized enrichment or reorganization of phosphorylated biomolecules, consistent with CBD’s known role in modulating protein phosphorylation pathways.[38],[39]

**Figure 5.**
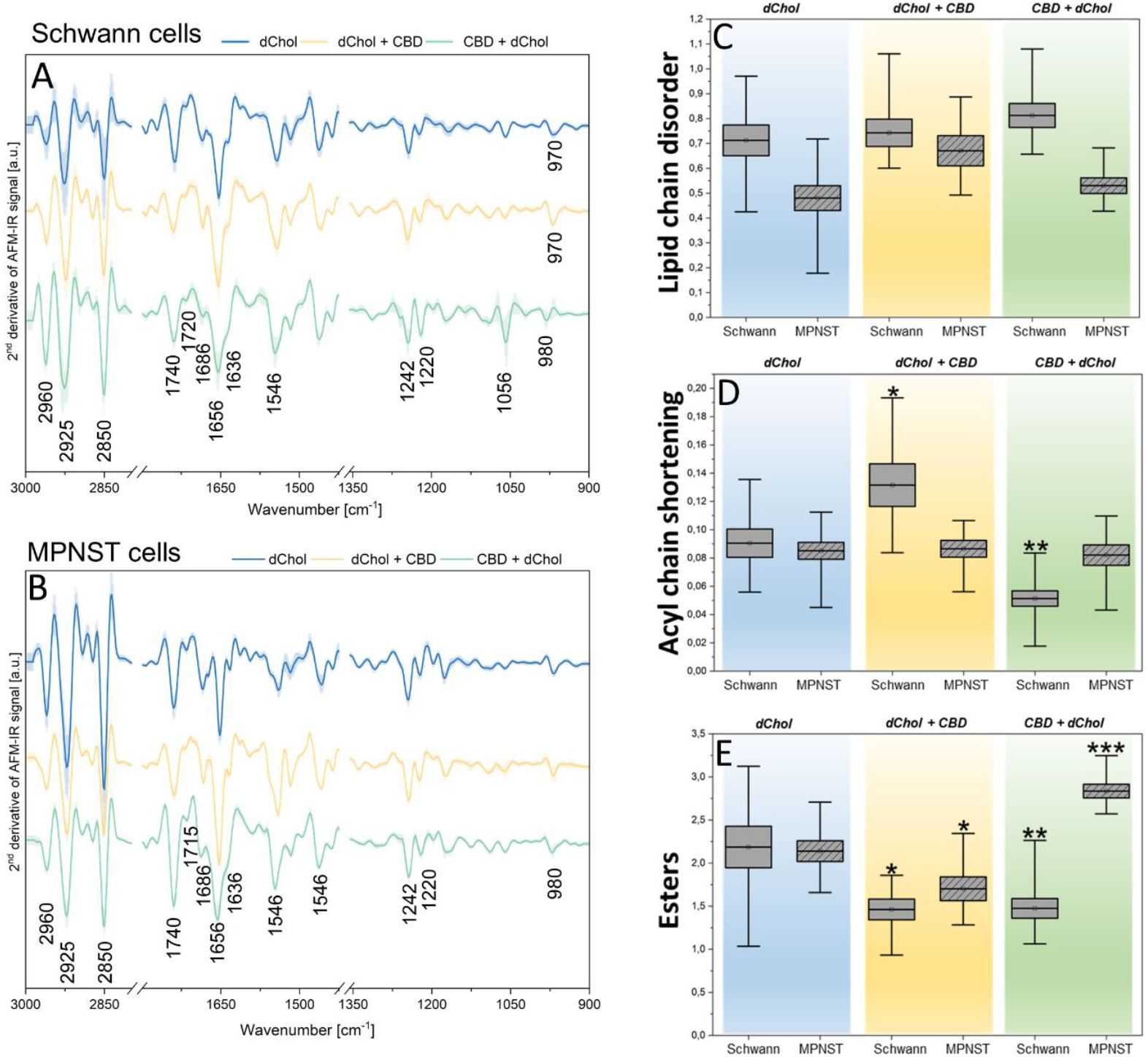
Second-derivative AFM-IR spectra of Schwann and MPNST cells (A, B) treated with 50µM d_6_-cholesterol (blue spectra), 50µM d_6_-cholesterol/3µM CBD (yellow spectra) and 3µM CBD/50µM d_6_-cholesterol (green spectra). Box diagrams (C-E) were constructed using integral intensity values calculated from second-derivative spectra collected across the entire cellular cross-section. Semi-quantitative evaluation allows assessment of lipid chain disorder (C: 2960 cm^-1^/2850 cm^-1^), acyl chain shortening (D:2870 cm^-1^/2925 cm^-1^) and ester content (E: 1740 cm^-1^). Statistical significance was determined using the one-way ANOVA (***p<0.001, p**<0.01, p*<0.05).

Integral analysis of selected AFM-IR bands enabled assessment of lipid-chain disorder, acyl-chain shortening, and esterified-lipid content (Fig. 5 C-E). For both cell lines, combined dChol/CBD treatment did not affect lipid-chain disorder, as reflected by an unchanged CH_3_/CH_2_ ratio.[40] CBD induced more pronounced alterations in lipid acyl chain organization in Schwann cells than in MPNST cells (Fig. 5, D). Supplementation with dChol followed by CBD led to increased acyl-chain shortening in Schwann cells (mean values: from 0.09 to 0.13), indicating a higher proportion of short-chain fatty acids (Fig. 5, D). This pattern aligns with enhanced lipid remodeling or β-oxidation, supporting the notion that CBD modulates intracellular lipid metabolism and LD composition.[7],[41] Conversely, when CBD was administered prior to dChol, the acyl-chain-shortening index decreased (mean values: from 0.09 to 0.05), suggesting enrichment in long-chain lipid species and potential suppression of fatty-acid shortening or β-oxidation, thereby promoting retention or synthesis of longer acyl chains.[42] None of these effects were observed in MPNST cells (Fig. 5, D). Esterified lipids, evaluated via the 1740 cm^−1^ band, exhibited further differences between cell types (Fig. 5, E). In Schwann cells, regardless of treatment order, esterified-lipid content decreased approximately 1.5-fold relative to baseline (mean values: from 2.19 to 1.46), possibly due to reduced fatty-acid availability required for cholesteryl-ester synthesis.[41] In MPNST cells, the probe-first treatment also reduced esterified lipids, whereas CBD-first exposure enhanced this fraction, indicating distinct or reprogrammed ester metabolism pathways in the malignant phenotype.

Beyond nanoscale chemical reorganization of lipid chains, AFM-IR imaging provided complementary insight into morphological changes within lipid-rich regions of the cytoplasm following CBD treatment. Cells pre-exposed to CBD prior to dChol supplementation exhibited heterogeneous populations of lipid-associated structures differing in size, shape, and spatial distribution (Fig. 6). Such morphological diversity is consistent with heterogeneous lipid storage states and may reflect different stages of LDs formation and remodeling. During the early stages of LDs formation, nascent vesicular structures became morphologically distinguishable within the cytoplasm (Fig. 6, A). These early lipid-associated features appeared as small, poorly defined vesicles, consistent with initial stages of LD biogenesis originating from the endoplasmic reticulum (ER).[43] At more advanced stages, distinct sharp crystalline structures were observed attached to the vesicular bodies (Fig. 6, B), indicating further lipid accumulation and reorganization. In addition, larger crystalline assemblies were detected at the cell surface and in adjacent extracellular regions (Fig. 6, C). These structures displayed well-defined morphologies with various orientations to the cell body.

**Figure 6.**
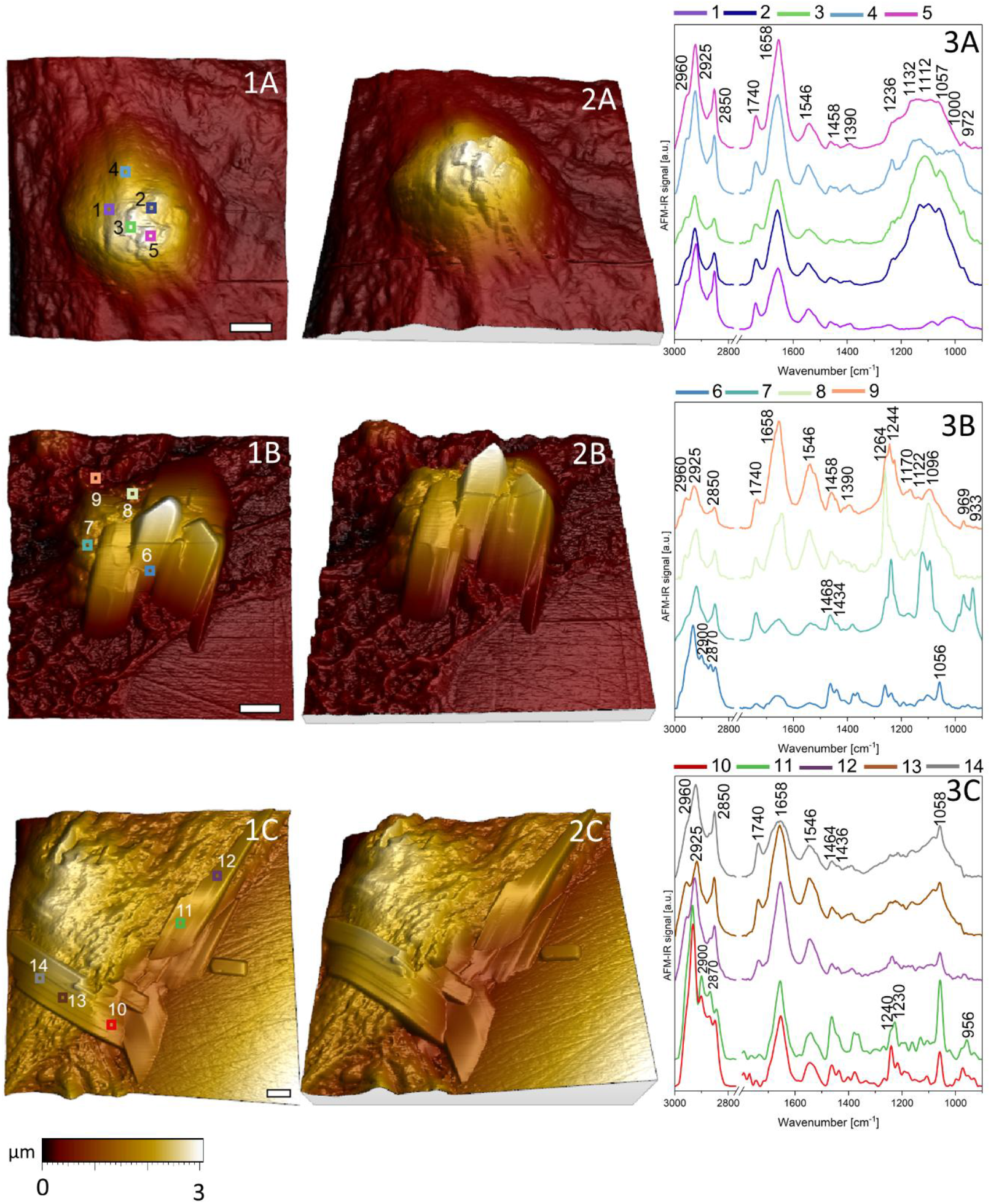
3D AFM visualization of morphological structures observed in peripheral glia cells after pre-incubation with 3 µM CBD followed by 50µM d_6_-cholesterol treatment (1A-2C). Local AFM-IR spectra correspond to the points and colors indicated on each topography map (3A-C). Scale bar equals 2µm.

AFM-IR spectra acquired from early-stage vesicular structures (Fig. 6, 3A) were dominated by vibrational features characteristic of ER-derived phospholipids. In addition to the ester carbonyl band at 1740 cm^−1^ and protein-related amide I and II bands at 1658 cm^−1^ and 1546 cm^−1^, respectively, a broad spectral region spanning 1236-972 cm^−1^ was observed. This region contained bands at 1236 cm^−1^, 1132-1112 cm^−1^, 1057 cm^−1^, 1000 cm^−1^, and 972 cm^−1^, corresponding to PO_2_^−^ stretching modes and C–O–P/C–O–C vibrations associated with glycerophospholipids, glycosidic linkages, and other phospholipid-or glycolipid-related components. [11],[44] Given that LDs originate from ER regions enriched in ribosomes and RNA, part of the spectral contribution within the 1236-972 cm^−1^ range likely arises from ribose- and phosphate-containing biomolecules localized around nascent droplets. [36],[45] Spectra collected from vesicles bearing crystalline attachments at later stages (Fig. 6, 3B) revealed clear lipid signatures. The presence of strong C–H stretching vibrations in the 2952-2850 cm^−1^ region indicated lipid-rich composition, attributable to cholesterol or its esterified forms, the latter supported by the appearance of a minor ester carbonyl band at 1740 cm^−1^ (Fig. S2, B). Vesicle-associated spectra further displayed bands at 1264-1244 cm^−1^ and a characteristic doublet at 1121 cm^−1^ and 1096 cm^−1^, consistent with phosphatidylcholine, in agreement with the phospholipid monolayer enveloping LDs during their formation. [46],[47] Surrounding cytoplasmic regions exhibited overlapping phospholipid-related features (spectra 8-9; 1260 cm^−1^ and 1244 cm^−1^), reflecting integration of LD-associated signals with the local cellular matrix. Spectral profiles of larger extracellular crystalline structures were dominated by cholesterol-specific vibrational signatures (Fig. 6, 3C; spectra 10-11; Fig. 1SI B). Variations in the relative intensities of characteristic bands (e.g., 1464 cm^−1^/1436 cm^−1^ and 1240 cm^−1^/1230 cm^−1^) depended on crystal orientation with respect to the cell, indicating anisotropic molecular organization. Additional spectra acquired from crystal-adjacent regions retained the characteristic cholesterol band at 1058 cm^−1^ superimposed on typical cellular spectral features (Fig. 6, 3C; spectra 12-14).

## Conclusions

This work introduces a novel multimodal spectrolipidomic sensing strategy that integrates Raman and FT-IR imaging, as well as AFM-IR nanospectroscopy with a deuterium-labeled cholesterol probe (dChol) for investigation of cellular lipid metabolism. By exploiting the intrinsic vibrational activity of dChol, this platform enables chemically selective, label-free tracking of cholesterol uptake and redistribution under CBD influence.

Raman spectroscopy emerged as the most effective technique for semi-quantitative detection of deuterated lipids in cells, achieving reliable identification of C–D stretching bands at concentrations as low as 10 µM-five times lower than the detection limit of conventional FT-IR imaging. These findings position Raman spectroscopy as a superior analytical readout for active deuterated lipid probes under native cellular conditions. At the nanoscale, beyond direct C–D vibrational detection, alternative lipid-associated marker was identified, with multivariate analysis revealing the ester carbonyl band (∼1740 cm^−1^) as a robust correlate of d_6_-cholesterol accumulation. AFM topographic mapping captured a concentration-dependent increase in lipid droplet related surface indentations, linking lipid accumulation to nanoscale deformation of the nuclear region.

Importantly, the combined spectroscopic approach demonstrated that depending on the sequence of CBD administration, it critically determines intracellular cholesterol handling. Raman and AFM-IR analyses consistently showed that CBD pre-exposure promotes enhanced cholesterol retention and lipid droplet remodeling, whereas CBD applied after probe supplementation restricts cholesterol accessibility. This highlights the ability of the presented framework to capture dynamic, sequence-dependent drug-lipid interactions. Moreover, this work demonstrates that AFM-IR can resolve successive stages of lipid droplet formation at the nanoscale by correlating morphological features with localized biochemical signatures.

Overall, this study establishes deuterium-labeled cholesterol as an active spectroscopic sensor and demonstrates that multimodal vibrational spectroscopy can extend molecular sensing beyond classical spectral markers to include nanoscale chemical and morphological readouts.

## CRediT authorship contribution statement

**Karolina Chrabąszcz:** conceptualization, data curation, formal analysis, funding acquisition, investigation, methodology, project administration, resources, supervision, validation, visualization, writing─original draft, writing─review and editing.

**Tycjan Kossowski-Kołodziej:** formal analysis, investigation, writing─review and editing

**Agnieszka Panek:** investigation, methodology, writing─review and editing

**Katarzyna Pogoda:** methodology, supervision, writing─review and editing

## Acknowledgments

This work was supported by the National Science Centre, Poland (2023/51/D/ST4/01686).

## Data Availability Statement

The data supporting the findings of this study are available in the RODBUK repository at https://doi.org/10.48733/IFJPAN/6IMDJA, including all datasets used to generate the figures presented in this manuscript.

## Appendix A. Supplementary data

Supplementary material related to this article can be found, in the online version, at doi:

## References

[1] D. Yang, X. Wang, L. Zhang, Y. Fang, Q. Zheng, X. Liu, W. Yu, S. Chen, J. Ying, F. Hua, Lipid metabolism and storage in neuroglia: role in brain development and neurodegenerative diseases, Cell Biosci. 12 (2022) 106. 10.1186/S13578-022-00828-0.

[2] R. Chrast, G. Saher, K.A. Nave, M.H.G. Verheijen, Lipid metabolism in myelinating glial cells: lessons from human inherited disorders and mouse models, J. Lipid Res. 52 (2011) 419. 10.1194/JLR.R009761.

[3] F.C.E. Vogel, A.B. Chaves-Filho, A. Schulze, Lipids as mediators of cancer progression and metastasis, Nat. Cancer 5 (2024) 16–29. 10.1038/S43018-023-00702-Z;SUBJMETA.

[4] S. Suppiah, S. Mansouri, Y. Mamatjan, J.C. Liu, M.M. Bhunia, V. Patil, P. Rath, B. Mehani, P. Heir, S. Bunda, G.L. Velez-Reyes, O. Singh, N. Ijad, N. Pirouzmand, T. Dalcourt, Y. Meng, S. Karimi, Q. Wei, F. Nassiri, T.J. Pugh, G.D. Bader, K.D. Aldape, D.A. Largaespada, G. Zadeh, Multiplatform molecular profiling uncovers two subgroups of malignant peripheral nerve sheath tumors with distinct therapeutic vulnerabilities, Nat. Commun. 2023 141 14 (2023) 1–15. 10.1038/s41467-023-38432-6.

[5] M. Tripson, K. Litwa, K. Soderstrom, Cannabidiol inhibits neuroinflammatory responses and circuit-associated synaptic loss following damage to a songbird vocal pre-motor cortical-like region, Sci. Reports 2023 131 13 (2023) 1–18. 10.1038/s41598-023-34924-z.

[6] S. Atalay, I. Jarocka-karpowicz, E. Skrzydlewskas, Antioxidative and anti-inflammatory properties of cannabidiol, Antioxidants 9 (2020) 1–20. 10.3390/antiox9010021.

[7] K. Konstantynowicz-Nowicka, K. Sztolsztener, A. Chabowski, E. Harasim-Symbor, Cannabidiol and sphingolipid metabolism: an unexplored link offering a novel therapeutic approach against high-fat diet-induced hepatic insulin resistance, J. Nutr. Biochem. 146 (2025) 109865. 10.1016/J.JNUTBIO.2025.109865.

[8] S.E. Guard, D.A. Chapnick, Z.C. Poss, C.C. Ebmeier, J. Jacobsen, T. Nemkov, K.A. Ball, K.J. Webb, H.L. Simpson, S. Coleman, E. Bunker, A. Ramirez, J.A. Reisz, R. Sievers, M.H.B. Stowell, A. D’Alessandro, X. Liu, W.M. Old, Multiomic Analysis Reveals Disruption of Cholesterol Homeostasis by Cannabidiol in Human Cell Lines, Mol. Cell. Proteomics 21 (2022) 100262. 10.1016/J.MCPRO.2022.100262.

[9] A.R. Watkins, Cannabinoid interactions with ion channels and receptors, Channels 13 (2019) 162–167. 10.1080/19336950.2019.1615824.

[10] E. Perez, J. Ceja-Vega, M. Krmic, A. Gamez Hernandez, J. Gudyka, R. Porteus, S. Lee, Differential Interaction of Cannabidiol with Biomembranes Dependent on Cholesterol Concentration, ACS Chem. Neurosci. 13 (2022) 1046–1054. 10.1021/ACSCHEMNEURO.2C00040/ASSET/IMAGES/LARGE/CN2C00040_0006.JPEG.

[11] K. Chrabąszcz, K. Pogoda, K. Cieżak, A. Panek, W.M. Kwiatek, Sensing Biomolecules Associated with Cells’ Radiosusceptibility by Advanced Micro- and Nanospectroscopy Techniques, ACS Sensors 9 (2024) 4887–4897. 10.1021/ACSSENSORS.4C01455/ASSET/IMAGES/LARGE/SE4C01455_0006.JPEG.

[12] E.S. Seltzer, A.K. Watters, D. Mackenzie, L.M. Granat, D. Zhang, Cannabidiol (Cbd) as a promising anti-cancer drug, Cancers (Basel). 12 (2020) 1–26. 10.3390/cancers12113203.

[13] X. Fu, Z. Yu, F. Fang, W. Zhou, Y. Bai, Z. Jiang, B. Yang, Y. Sun, X. Tian, G. Liu, Cannabidiol attenuates lipid metabolism and induces CB1 receptor-mediated ER stress associated apoptosis in ovarian cancer cells, Sci. Reports 2025 151 15 (2025) 1–16. 10.1038/s41598-025-88917-1.

[14] N. Mangal, S. Erridge, N. Habib, A. Sadanandam, V. Reebye, M.H. Sodergren, Cannabinoids in the landscape of cancer, J. Cancer Res. Clin. Oncol. 147 (2021) 2507– 2534. 10.1007/S00432-021-03710-7.

[15] C. De La Haba, J.R. Palacio, P. Martínez, A. Morros, Effect of oxidative stress on plasma membrane fluidity of THP-1 induced macrophages, Biochim. Biophys. Acta - Biomembr. 1828 (2013) 357–364. 10.1016/J.BBAMEM.2012.08.013.

[16] S. Li, H. Yuan, L. Li, Q. Li, P. Lin, K. Li, Oxidative Stress and Reprogramming of Lipid Metabolism in Cancers, Antioxidants 2025, Vol. 14, Page 201 14 (2025) 201. 10.3390/ANTIOX14020201.

[17] L. Tirinato, M.G. Marafioti, F. Pagliari, J. Jansen, I. Aversa, R. Hanley, C. Nisticò, D. Garcia-Calderón, G. Genard, J.F. Guerreiro, F.S. Costanzo, J. Seco, Lipid droplets and ferritin heavy chain: a devilish liaison in human cancer cell radioresistance, Elife 10 (2021). 10.7554/ELIFE.72943.

[18] A.J. Hoy, S.R. Nagarajan, L.M. Butler, Tumour fatty acid metabolism in the context of therapy resistance and obesity, Nat. Rev. Cancer 21 (2021) 753–766. 10.1038/S41568-021-00388-4,.

[19] X. Huang, Z. Xue, D. Zhang, H.J. Lee, Pinpointing Fat Molecules: Advances in Coherent Raman Scattering Microscopy for Lipid Metabolism, Anal. Chem. 96 (2024) 7945–7958. 10.1021/ACS.ANALCHEM.4C01398.

[20] K. Majzner, K. Kochan, N. Kachamakova-Trojanowska, E. Maslak, S. Chlopicki, M. Baranska, Raman imaging providing insights into chemical composition of lipid droplets of different size and origin: in hepatocytes and endothelium, Anal. Chem. 86 (2014) 6666–6674. 10.1021/AC501395G.

[21] Y. Bai, C.M. Camargo, S.M.K. Glasauer, R. Gifford, X. Tian, A.P. Longhini, K.S. Kosik, Single-cell mapping of lipid metabolites using an infrared probe in human-derived model systems, Nat. Commun. 2024 151 15 (2024) 1–15. 10.1038/s41467-023-44675-0.

[22] A. Alfonso-García, S.G. Pfisterer, H. Riezman, E. Ikonen, E.O. Potma, D38-cholesterol as a Raman active probe for imaging intracellular cholesterol storage, J. Biomed. Opt. 21 (2016) 061003. 10.1117/1.JBO.21.6.061003.

[23] S. Egoshi, K. Dodo, M. Sodeoka, Deuterium Raman imaging for lipid analysis, Curr. Opin. Chem. Biol. 70 (2022) 102181. 10.1016/J.CBPA.2022.102181.

[24] A.N. Omelchenko, K.A. Okotrub, T.N. Igonina, T.A. Rakhmanova, S. V. Okotrub, I.N. Rozhkova, V.S. Kozeneva, S.Y. Amstislavsky, N. V. Surovtsev, Probing metabolism in mouse embryos using Raman spectroscopy and deuterium tags, Spectrochim. Acta Part A Mol. Biomol. Spectrosc. 325 (2025) 125044. 10.1016/J.SAA.2024.125044.

[25] S.O. Shuster, A.E. Curtis, C.M. Davis, Optical Photothermal Infrared Imaging Using Metabolic Probes in Biological Systems, Anal. Chem. 97 (2025) 8202–8212. 10.1021/ACS.ANALCHEM.4C03752.

[26] K. Chrab, N. Piergies, A. Panek, M. Szczepanek-dulska, K. Pogoda, Toward precision in nanoscale IR spectroscopy: Optimizing biological sample preparation for molecular and morphological integrity, 350 (2026).

[27] N. Piergies, J. Mathurin, A. Dazzi, A. Deniset-Besseau, M. Oćwieja, C. Paluszkiewicz, W.M. Kwiatek, IR nanospectroscopy to decipher drug/metal nanoparticle interactions: Towards a better understanding of the spectral signal enhancement and its distribution, Appl. Surf. Sci. 609 (2023) 155217. 10.1016/J.APSUSC.2022.155217.

[28] J. Yang, C. Shay, N.F. Saba, Y. Teng, Cancer metabolism and carcinogenesis, Exp. Hematol. Oncol. 2024 131 13 (2024) 1–14. 10.1186/S40164-024-00482-X.

[29] L.A. Broadfield, A.A. Pane, A. Talebi, J. V. Swinnen, S.M. Fendt, Lipid metabolism in cancer: New perspectives and emerging mechanisms, Dev. Cell 56 (2021) 1363–1393. 10.1016/J.DEVCEL.2021.04.013.

[30] P.B. Jonker, A. Muir, Metabolic ripple effects – deciphering how lipid metabolism in cancer interfaces with the tumor microenvironment, Dis. Model. Mech. 17 (2024) dmm050814. 10.1242/DMM.050814.

[31] A.E. Loneker, F. Alisafaei, A. Kant, D. Li, P.A. Janmey, V.B. Shenoy, R.G. Wells, Lipid droplets are intracellular mechanical stressors that impair hepatocyte function, Proc. Natl. Acad. Sci. U. S. A. 120 (2023) e2216811120. 10.1073/PNAS.2216811120.

[32] Y. Li, P. Khanal, F. Norheim, M. Hjorth, T. Bjellaas, C.A. Drevon, J. Vaage, A.R. Kimmel, K.T. Dalen, Plin2 deletion increases cholesteryl ester lipid droplet content and disturbs cholesterol balance in adrenal cortex, J. Lipid Res. 62 (2021). 10.1016/J.JLR.2021.100048.

[33] F. Les, M. Sofía Valero, M.P. Arruebo, A. Wró Nski, I. Dobrzy’nska, D. Dobrzy’nska, S. Ş Ekowski, W. Łuczaj, E. Olchowik-Grabarek, E. Zbieta Skrzydlewska, Cannabidiol and Cannabigerol Modify the Composition and Physicochemical Properties of Keratinocyte Membranes Exposed to UVA, Int. J. Mol. Sci. 2023, Vol. 24, Page 12424 24 (2023) 12424. 10.3390/IJMS241512424.

[34] C.L.E. Yen, S.J. Stone, S. Koliwad, C. Harris, R. V. Farese, Thematic Review Series: Glycerolipids. DGAT enzymes and triacylglycerol biosynthesis, J. Lipid Res. 49 (2008) 2283–2301. 10.1194/JLR.R800018-JLR200.

[35] N. Rimmerman, A. Juknat, E. Kozela, R. Levy, H.B. Bradshaw, Z. Vogel, The non-psychoactive plant cannabinoid, cannabidiol affects cholesterol metabolism-related genes in microglial cells, Cell. Mol. Neurobiol. 31 (2011) 921–930. 10.1007/S10571-011-9692-3.

[36] P. Gardner, A. Spadea, J. Denbigh, M.J. Lawrence, M. Kansiz, Analysis of fixed and live single cells using optical photothermal infrared with concomitant Raman spectroscopy, Anal. Chem. 93 (2021) 3938–3950. 10.1021/ACS.ANALCHEM.0C04846/ASSET/IMAGES/LARGE/AC0C04846_0008.JPEG.

[37] S. Alali, G. Riazi, M.R. Ashrafi-Kooshk, S. Meknatkhah, S. Ahmadian, M.H. Ardakani, B. Hosseinkhani, Cannabidiol Inhibits Tau Aggregation In Vitro, Cells 10 (2021). 10.3390/CELLS10123521.

[38] M.N.M. Volmar, J. Cheng, H. Alenezi, S. Richter, A. Haug, Z. Hassan, M. Goldberg, Y. Li, M. Hou, C. Herold-Mende, C.L. Maire, K. Lamszus, C. Flüh, J. Held-Feindt, G. Gargiulo, G.J. Topping, F. Schilling, Di. Saur, G. Schneider, M. Synowitz, J.A. Schick, R.E. Kälin, R. Glass, Cannabidiol converts NF-κB into a tumor suppressor in glioblastoma with defined antioxidative properties, Neuro. Oncol. 23 (2021) 1898– 1910. 10.1093/NEUONC/NOAB095.

[39] T.A.M. Vrechi, A.H.F.F. Leão, I.B.M. Morais, V.C. Abílio, A.W. Zuardi, J.E.C. Hallak, J.A. Crippa, C. Bincoletto, R.P. Ureshino, S.S. Smaili, G.J.S. Pereira, Cannabidiol induces autophagy via ERK1/2 activation in neural cells, Sci. Rep. 11 (2021) 5434. 10.1038/S41598-021-84879-2.

[40] A. Blat, J. Dybas, M. Kaczmarska, K. Chrabaszcz, K. Bulat, R.B. Kostogrys, A. Cernescu, K. Malek, K.M. Marzec, An analysis of isolated and intact rbc membranes - a comparison of a semiquantitative approach by means of FTIR, Nano-FTIR, and Raman Spectroscopies, Anal. Chem. 91 (2019). 10.1021/acs.analchem.9b01536.

[41] P. Bielawiec, S. Dziemitko, K. Konstantynowicz-Nowicka, A. Chabowski, J. Dzięcioł, E. Harasim-Symbor, Cannabidiol improves muscular lipid profile by affecting the expression of fatty acid transporters and inhibiting de novo lipogenesis, Sci. Rep. 13 (2023) 1–16. 10.1038/s41598-023-30872-w.

[42] E. Harasim-Symbor, P. Bielawiec, A. Pedzinska-Betiuk, J. Weresa, B. Malinowska, K. Konstantynowicz-Nowicka, A. Chabowski, Cannabidiol treatment changes myocardial lipid profile in spontaneously hypertensive rats, Nutr. Metab. Cardiovasc. Dis. 34 (2024) 2817–2833. 10.1016/J.NUMECD.2023.07.007.

[43] R. Dhiman, R.S. Perera, C.S. Poojari, H.T.A. Wiedemann, R. Kappl, C.W.M. Kay, J.S. Hub, B. Schrul, Hairpin protein partitioning from the ER to lipid droplets involves major structural rearrangements, Nat. Commun. 2024 151 15 (2024) 4504-. 10.1038/s41467-024-48843-8.

[44] A. Blat, J. Dybas, M. Kaczmarska, K. Chrabaszcz, K. Bulat, R.B. Kostogrys, A. Cernescu, K. Malek, K.M. Marzec, An analysis of isolated and intact rbc membranes - a comparison of a semiquantitative approach by means of FTIR, Nano-FTIR, and Raman Spectroscopies, Anal. Chem. 91 (2019) 9867–9874. 10.1021/ACS.ANALCHEM.9B01536/SUPPL_FILE/AC9B01536_SI_001.PDF.

[45] S. Cottier, R. Schneiter, Lipid droplets form a network interconnected by the endoplasmic reticulum through which their proteins equilibrate, J. Cell Sci. 135 (2022). 10.1242/JCS.258819.

[46] T.C. Walther, J. Chung, R. V. Farese, Lipid Droplet Biogenesis, Annu. Rev. Cell Dev. Biol. 33 (2017) 491–510. 10.1146/ANNUREV-CELLBIO-100616-060608.

[47] K. Chrabąszcz, Spectroscopic active probes for investigation of lipids transformation in cells and membranes, J. Phys. Chem. B (n.d.). 10.1021/acs.jpcb.5c05981.

