## Supporting Information for "Spectrolipidomics of glial cell lines: a deuterated probe for semiquantitative monitoring of cannabidiol-induced cholesterol modulation"

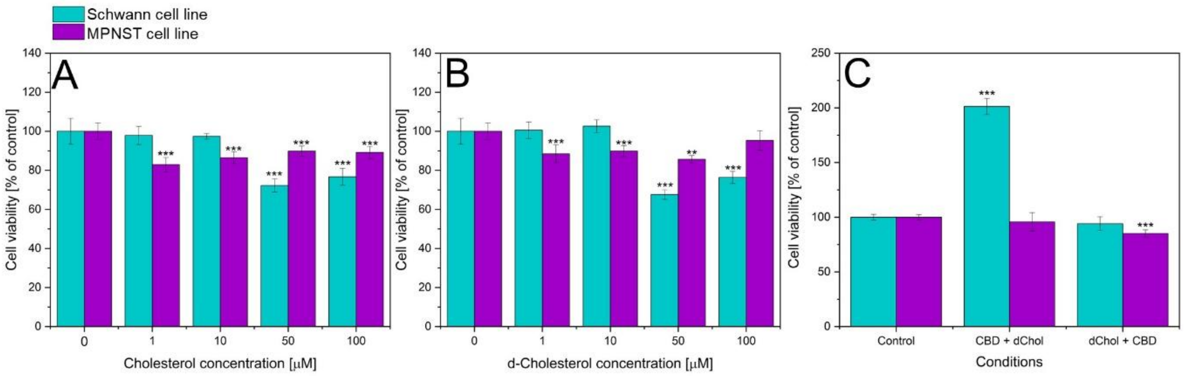


**Figure S1.** Results of the MTS assay for Schwann (turquoise) and MPNST (violet) cell lines: (A) different concentrations of cholesterol, (B) $d_6$-cholesterol, and (C) mixtures of 3 μM CBD with 50 μM $d_6$-cholesterol (***p<0.001, p**<0.01).

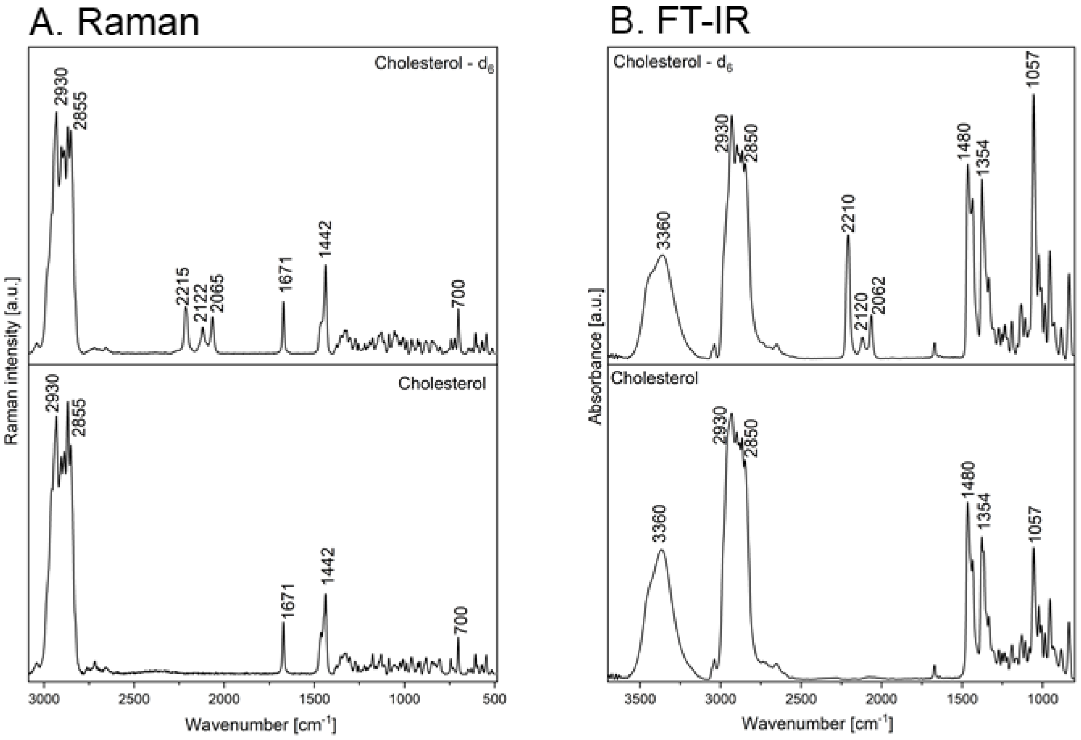


**Figure S2.** Comparison of cholesterol and $d_6$-cholesterol spectra acquired using (A) Raman and (B) FT-IR spectroscopy.

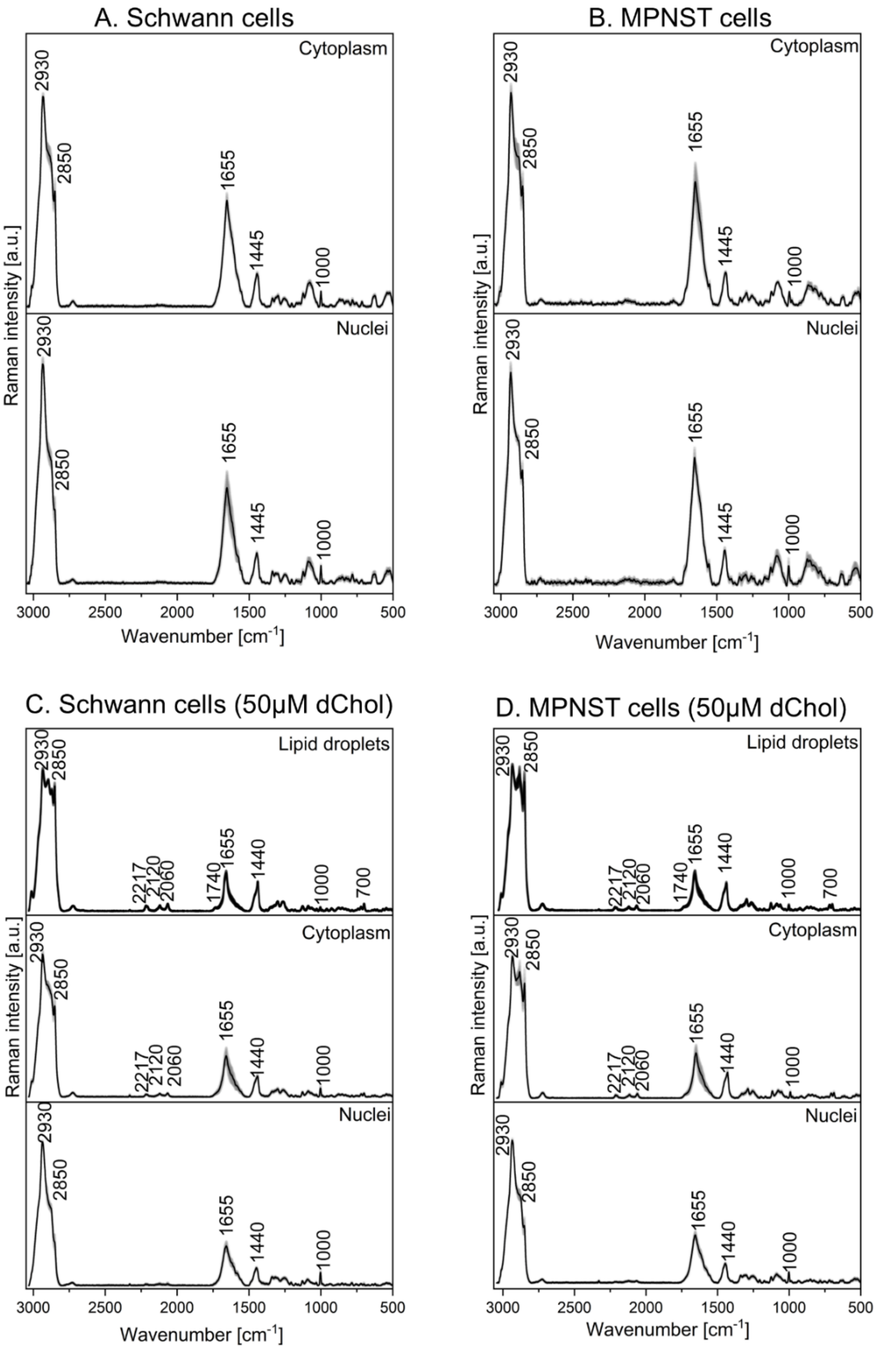


**Figure S3.** Mean Raman spectra of cytoplasm and nuclei obtained from single-cell cluster analysis for (A) Schwann and (B) MPNST cells. Following treatment with 50 µM $d_6$-cholesterol, in addition to the differentiation between cytoplasm and nuclei, lipid droplets containing visible C–D vibrational bands from the deuterated probe were also identified (C, D).

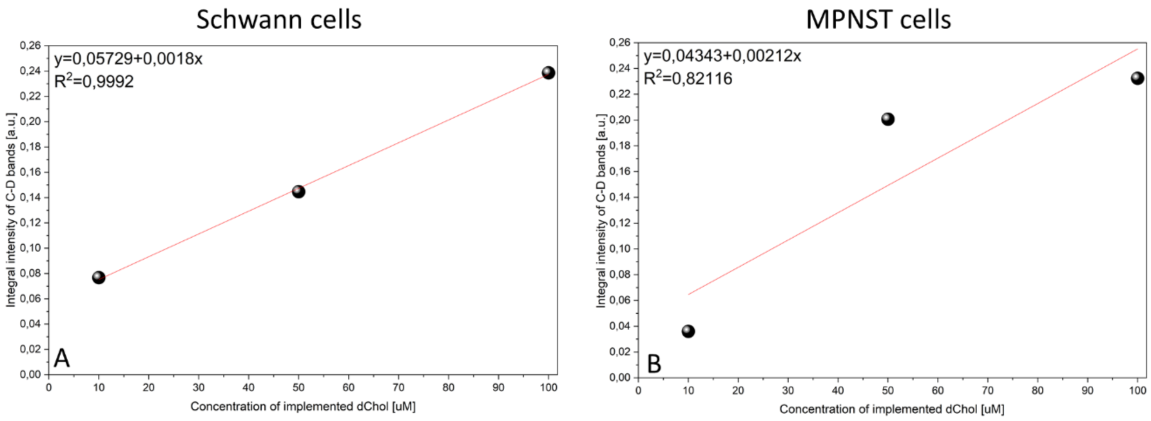


**Figure S4.** Linear dependence between the applied dChol concentrations and the integrated C–D band intensity in (A) Schwann and (B) MPNST cell lines.

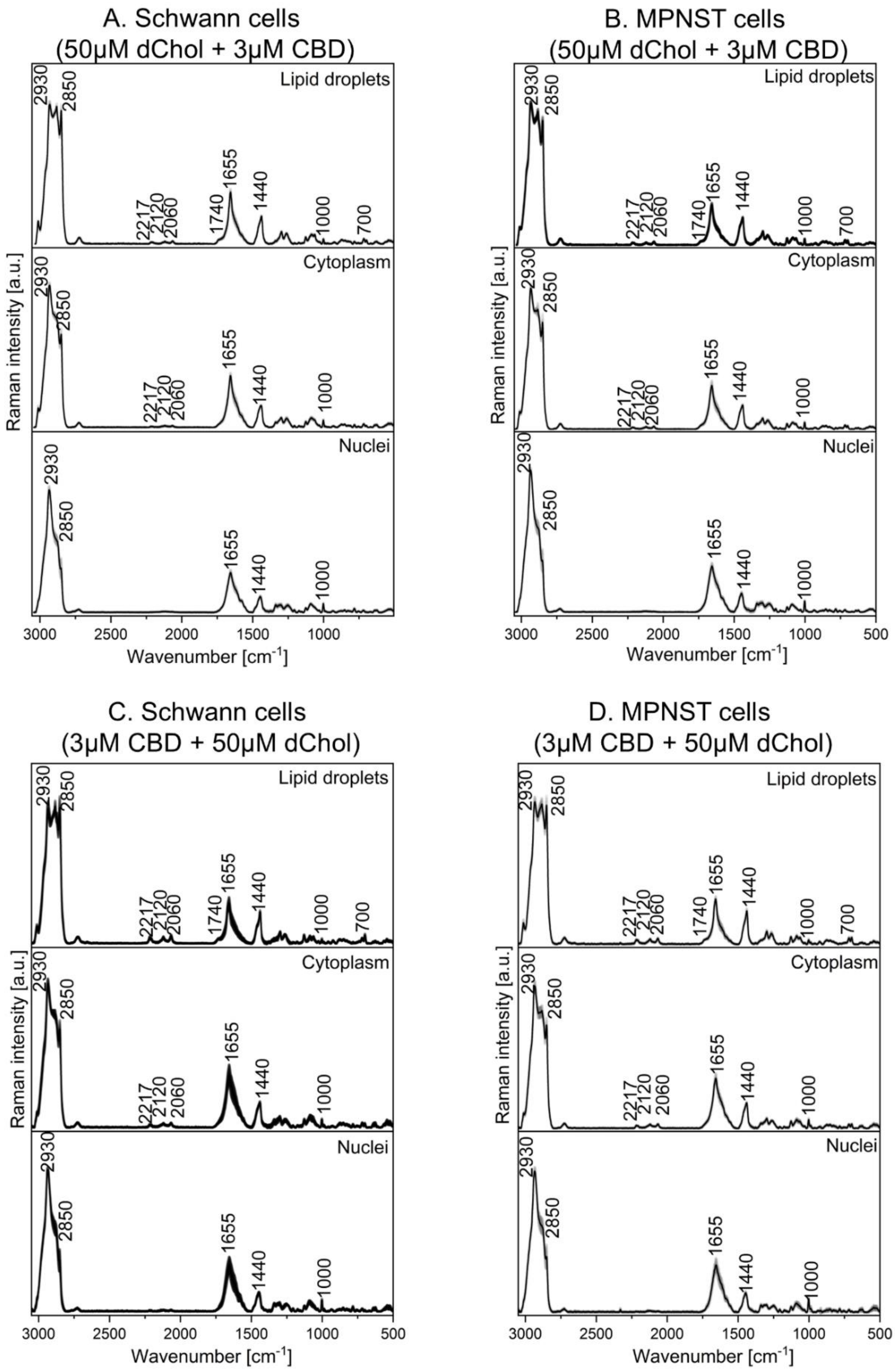


**Figure S5.** Mean Raman spectra of lipid droplets, cytoplasm, and nuclei obtained from single-cell cluster analysis of treated Schwann and MPNST cells. Spectral variations following treatment with (A, B) 50 μM $d_6$-cholesterol/3 μM CBD and (C, D) 3 μM CBD/50 μM $d_6$-cholesterol are presented.

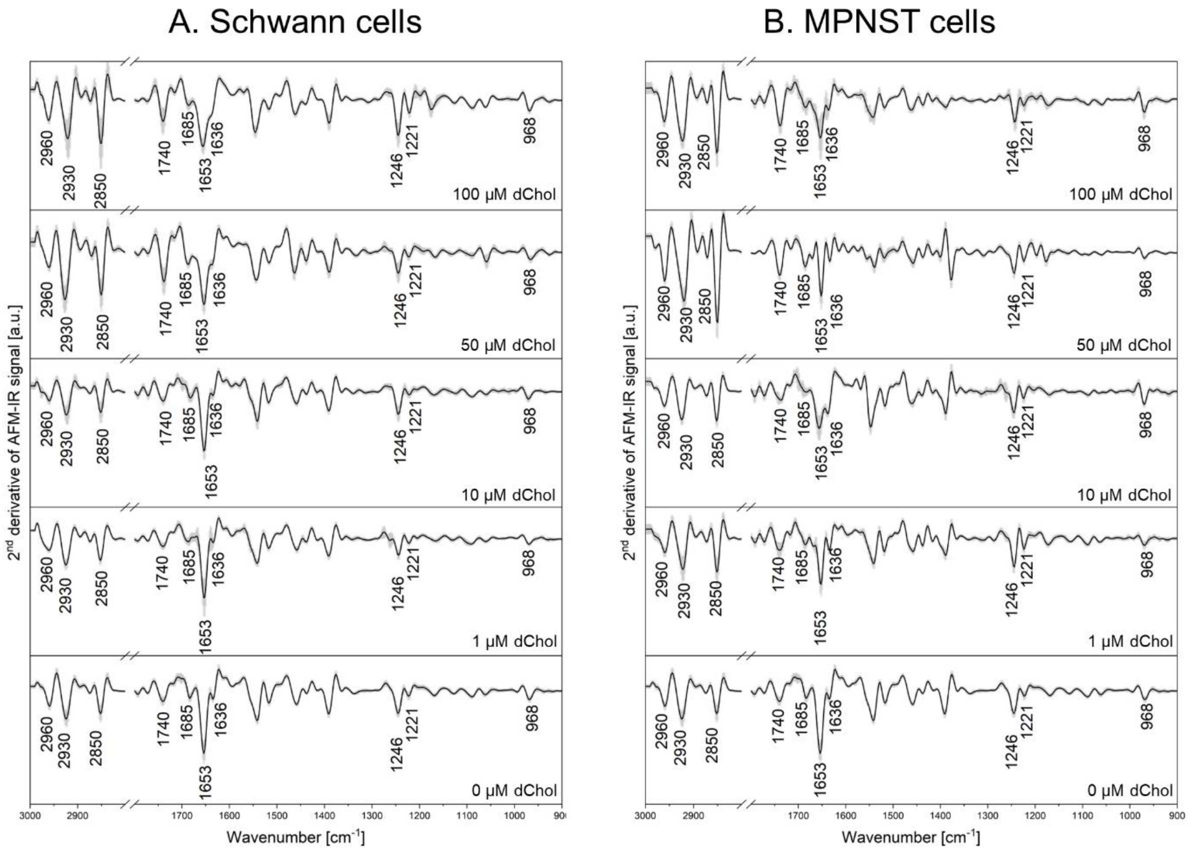


**Figure S6.** Comparison of second-derivative AFM-IR spectra collected for (A) Schwann and (B) MPNST cell lines treated with 1 µM, 10 µM, 50 µM, and 100 µM $d_6$-cholesterol.
